# Model based control can give rise to devaluation insensitive choice

**DOI:** 10.1101/2022.08.21.504635

**Authors:** Neil Garrett, Sean Allan, Nathaniel D. Daw

**Affiliations:** School of Psychology, University of East Anglia; Princeton Neuroscience Institute and Department of Psychology, Princeton University

## Abstract

Influential recent work aims to ground psychiatric dysfunction in the brain’s basic computational mechanisms. For instance, compulsive symptoms as in drug abuse have been argued to arise from imbalance between multiple systems for instrumental learning. Computational models suggest that such multiplicity arises because the brain adaptively simplifies laborious “model-based” deliberation by sometimes relying on a cheaper, more habitual “model-free” shortcut. Support for this account comes in part from failures to appropriately change behavior in light of new events. Notably, instrumental responding can, in some circumstances, persist despite reinforcer devaluation, perhaps reflecting control by model-free mechanisms that are driven by past reinforcement rather than knowledge of the (now devalued) outcome. However, another important line of theory – heretofore mostly studied in Pavlovian conditioning – posits a different mechanism that can also modulate behavioral change. It concerns how animals identify different rules or contingencies that may apply in different circumstances, by covertly clustering experiences into distinct groups identified with different “latent causes” or contexts. Such clustering has been used to explain the return of Pavlovian responding following extinction.

Here we combine both lines of theory to investigate the consequences of latent cause inference on instrumental sensitivity to reinforcer devaluation. We show that because segregating events into different latent clusters prevents generalization between them, instrumental insensitivity to reinforcer devaluation can arise in this theory even using only model-based planning, and does not require or imply any habitual, model-free component. In simulations, these ersatz habits (like laboratory ones) emerge after overtraining, interact with contextual cues, and show preserved sensitivity to reinforcer devaluation on a separate consumption test, a standard control. While these results do not rule out a contribution of model-free learning *per se*, they point to a subtle and important role of state inference in instrumental learning and highlight the need for caution in using reinforcer devaluation procedures to rule in (or out) the contribution of different learning mechanisms. They also offer a new perspective on the neurocomputational substrates of drug abuse and the relevance of laboratory reinforcer devaluation procedures to this phenomenon.

## Introduction

A key idea across psychological and neural theories is that the brain judiciously simplifies laborious computations using heuristics or shortcuts (Daw et al., 2011; Tversky and Kahneman, 1974). One well-developed version of this idea concerns the tradeoff between deliberative and automatic modes of control, as operationalized in rodents using a widely studied reinforcer devaluation procedure (Adams and Dickinson, 1981; Colwill and Rescorla, 1985; Dickinson, 1985; Dickinson and Balleine, 2002; Rescorla, 1990) (**Figure 1a**). Here, animals are trained to leverpress for food, then tested following reinforcer devaluation (e.g., by taste aversion conditioning: pairing the food with illness). In some circumstances, such as when overtrained, animals nevertheless work persistently for the devalued outcome (**Figure 1c**). This failure to appropriately adjust behavior following reinforcer devaluation is widely viewed as a laboratory model of the familiar human experience of habits, whereby with repetition, some actions (such as making a particular turn on the way to work) seem to become automatized and we tend to slip and perform them even when contextually inappropriate (e.g., when we actually intend to go elsewhere). Pathological dominance of these same habit mechanisms has also been argued to produce the compulsive, seemingly consequence-insensitive behaviors that characterize disorders of compulsion (Gillan et al., 2016; Graybiel and Rauch, 2000) such as drug abuse (Everitt and Robbins, 2016, 2005).

**Figure 1.**
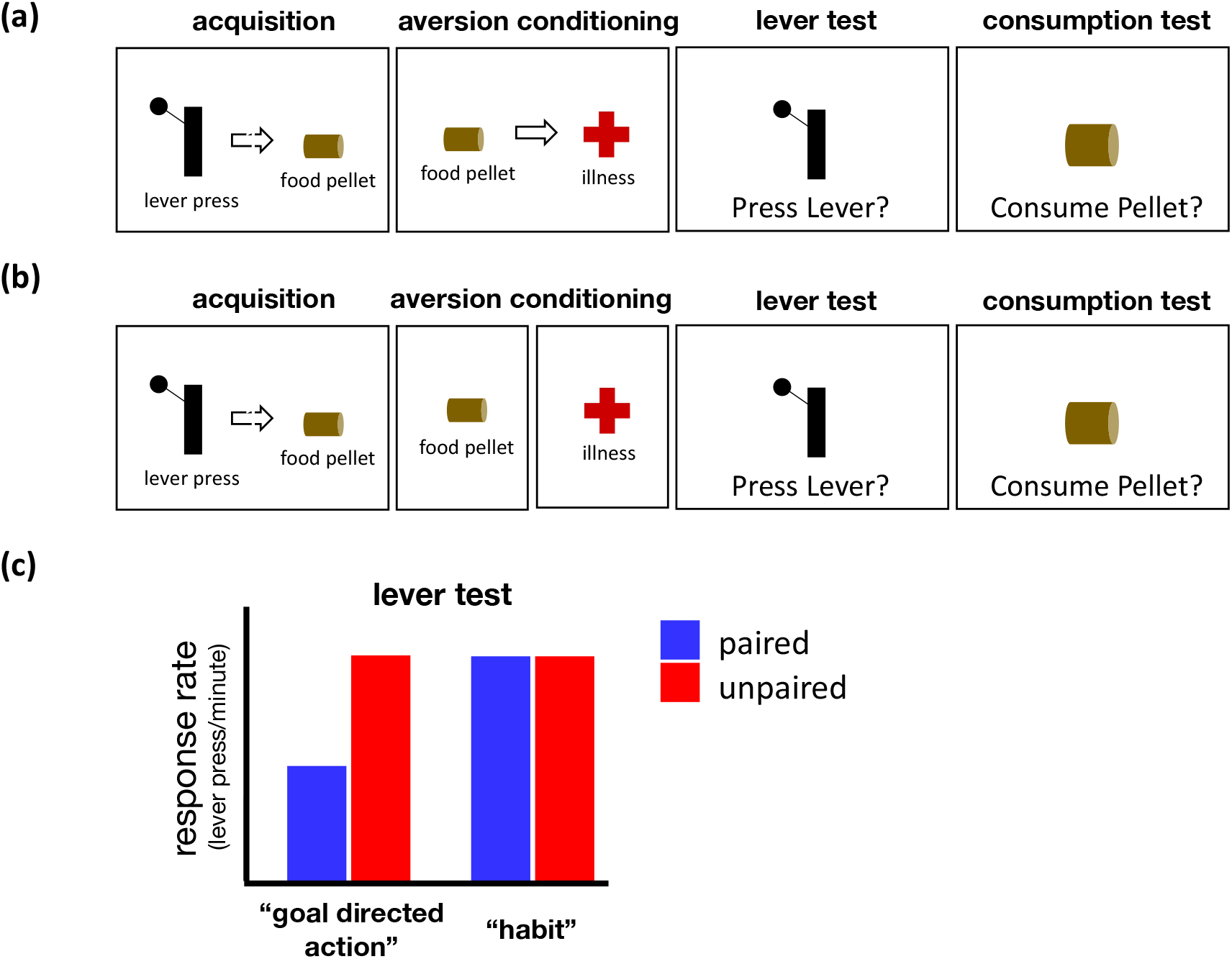
**(a)** Paired Condition. Typical experimental timeline used to test habit formation in which illness is paired with food during aversion conditioning which we base our simulations on. Animals are trained to leverpress for food (*acquisition*) then tested following *aversion conditioning* (pairing the food with illness, induced for example via injection of lithium chloride). A *consumption test* – in which the animal is freely provided with food pellets (without a lever present) – verifies the efficacy of the aversion conditioning. **(b)** Unpaired (control) Condition. Food and illness are separated during aversion conditioning preventing the formation of an association between the two. **(c)** In some circumstances, instrumental responding persists despite reinforcer devaluation (stylized data illustrated), a behavior which is viewed as arising from a reliance on habits (model free learning). This occurs, for example, when animals are overtrained (Holland, 2004) as well as when the context in which the aversion conditioning occurs differs to that used for the acquisition and test phases (Bouton et al., 2021).

An influential computational analysis, in turn, views these habitual behaviors as reflecting a simplified algorithmic strategy for evaluating candidate actions to decide what to do (Daw et al., 2005; Sutton and Barto, 2018). Although it is generally most accurate to use a learned “internal model” of the task contingencies iteratively to anticipate and evaluate an action’s consequences (a leverpress leads to food, which may or may not be desirable), such computation requires many steps which cheaper “model-free” methods may skip by simply storing the endpoint of this computation (e.g., the decision to leverpress, called a policy in reinforcement learning or a stimulus-response association in classic associative learning accounts). On this interpretation, the transition to habits reflects the brain shifting from laborious but accurate model-based planning to cheaper but approximate model-free responding. Because – for highly practiced behaviors in stable circumstances – this shortcut generally produces the same result (a leverpress) with less computation at choice time, this analysis justifies habits as reflecting a rational, circumstantially appropriate tradeoff between the costs of computation and the costs of error (Daw et al., 2005; Keramati et al., 2011).

However, a second and mostly separate line of influential theories details a different mechanism that may also contribute to changing – or, crucially, failing to change – behavior in light of new experience. Statistical accounts view Pavlovian conditioning as reflecting a process of inferring the statistical structure of events. In particular, these *latent cause inference* theories (Courville et al., 2004, 2003; Gershman et al., 2015, 2015, 2010) view the brain as adaptively clustering experiences into groups, representing different types of trials or different (“latent” or subjectively inferred) contexts in which different contingencies manifest. The rationale for these models is that experiences are drawn from different contingencies in different situations; and therefore learning requires, in part, figuring out which contingencies apply when. In effect, such clustering gates generalization: learning about contingencies applies within each context, but not between them. One particularly important application of this logic is in regard to the *extinction* of previously conditioned associations: specifically, findings that Pavlovian responding recurs even following extinction. By inferring that extinction trials in which a conditioned stimulus (CS) is no longer reinforced arise from a different latent context than did the initial acquisition trials, these theories explain many phenomena of renewal (Bouton and Bolles, 1979) and recovery (Pavlov, 1927) which imply that extinction learning coexists alongside initial acquisition learning, rather than simply erasing it (Gershman et al., 2010).

Though these theories have primarily been applied to Pavlovian conditioning, such latent grouping of contingencies into contexts is, in principle, equally relevant to instrumental learning. That is, the basic insight of these models applies to instrumental choice: that different task contingencies may occur in different circumstances, so the organism must simultaneously figure out which tasks are active while learning to perform them. Indeed, Schwöbel et al. (Schwöbel et al., 2021) recently put forward a theory nesting dual-process instrumental control (model-based learning alongside a modified model-free policy learner) underneath latent cause inference, and used it to simulate several results involving the making and breaking of habits.

Here we dive more deeply into the merger of these two lines of theory, by examining the implications of latent cause inference for fully deliberative model-based control alone – with a context-dependent learned world model but importantly without any model-free value or policy caching component. We show that these mechanisms alone can reproduce the devaluation-insensitive instrumental responding thought to be characteristic of habits, for reasons entirely analogous to why latent cause models explain failures of extinction. The account explains the characteristic emergence of habits with overtraining, as well as recent results concerning the effect of manipulations of training context on the reinforcer devaluation effect, which are on this view analogous to similar experiments exploring the context specificity of Pavlovian extinction. In particular, if the taste aversion conditioning used for reinforcer devaluation is attributed to a different latent context than the instrumental probe (a lever test), then even fully deliberative model-based control will exhibit instrumental insensitivity to reinforcer devaluation. Surprisingly, in simulations, we observe that taste aversion conditioning can generalise to an outcome consumption test (intended to verify taste aversion efficacy) despite also failing to generalise to the instrumental (lever) test, a pattern of results previously interpreted as ruling out model-based control.

In short, our model and simulation results demonstrate that instrumental insensitivity to reinforcer devaluation need not imply or reflect computational simplification such as model-free learning or stimulus-response habits. Conversely, the demonstration that insensitivity of action choice to outcome value can arise for entirely distinct computational reasons (involving contextual inference) offers a distinct formal perspective on what types of dysfunctional computations might contribute to pathologically consequence-insensitive choice, as in drug abuse, and a new interpretation of specific experimental evidence (often using a reinforcer devaluation procedure and related paradigms) purportedly tying habits to disorders. This new perspective naturally accommodates aspects of drug abuse (such as the existence and context sensitivity of craving, goal-directed drug-seeking, and relapse effects) that are hard to explain from a stimulus-response view alone.

## The theory: instrumental learning with latent contexts

We augment a standard theory of model-based instrumental learning – inferring an unknown Markov decision process (MDP) – with the possibility that different MDPs obtain on different trials. Learning in this setting – inference given the model – thus, roughly, nests model-based MDP learning and planning (Daw et al., 2005) under latent cause inference (Gershman et al., 2015, 2010) about which MDP is active.

### Generative model

To model conditioning as Bayesian inference, we first describe the statistical generative model that is assumed to govern task events. The agent then infers the task contingencies through standard inference in this model, and makes action choices appropriate to the inferred task. For the generative model, we assume an infinite mixture model over episodic, fully observable Markov decision processes. That is, for each trial, a latent cause (henceforth, “context”) is drawn that is associated with a particular MDP, which governs the resulting episode until termination. Although the states of each MDP are observable, the different MDPs can share the same states, so their occurrence can be aliased across different latent contexts. That is (unlike in the more usual fully observable setting) the agent must infer whether a particular state like the option to press a lever (S1 in **Figure 2a** and **2b**) is occurring in the same or different context as other previous experiences with a similar situation.

**Figure 2.**
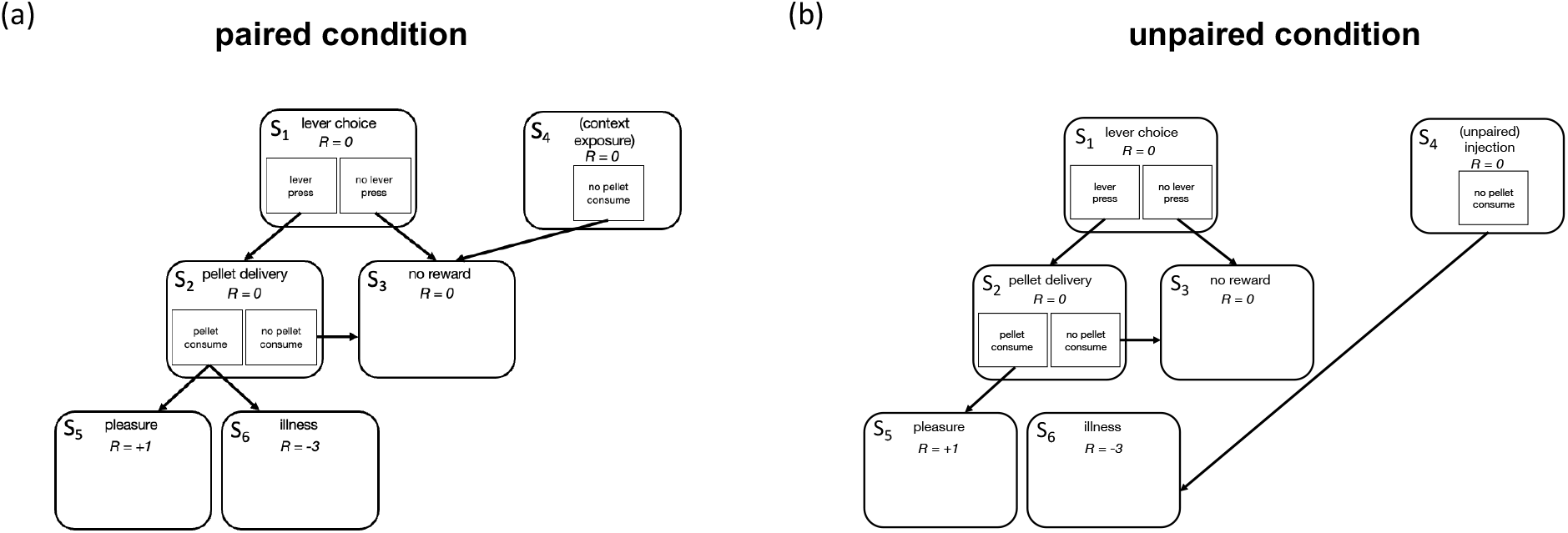
Task representation (MDP) of reward devaluation as represented by a goal directed system and used in our simulations. S_1_–S_6_ are the six possible states within the task. R={−3, 0, 1} represents rewards/losses obtained in each state. The agent can start in S_1_, S_2_ or S_4_, depending on the phase of the task. Each trial of instrumental leverpressing acquisition starts in S_1_ and proceeds to a rewarded outcome S_5_ given the appropriate choices. Taste aversion conditioning (a, for the paired condition) starts in S_2_ with the action “pellet consume” now transitioning to illness S_6_. In the control (unpaired) condition (b), animals instead start in S_4_ during their taste aversion condition and thus transition to illness S_6_ without encountering a food pellet S_2_. For counterbalancing the other group’s taste aversion conditioning exposure, animals in the paired condition also encounter trials that start in S_4_ but end in neutral outcome S_3_, while animals in the unpaired condition are exposed to food S_2_ while still transitioning to the same positive outcome S_5_ encountered during training.

More specifically, at each trial *t* a latent context *c*_*t*_ is drawn from an infinite multinomial mixture model according to a Chinese restaurant process prior, i.e., *c*_1_ = 1; 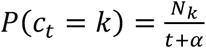 for a previously encountered context *k*, and 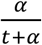 for a previously unobserved context. Here *N*_*k*_ is the number of times context *k* has previously occurred, and *α* is a concentration parameter which governs how often new causes are likely to be encountered. We used a very low value of *α*, 1*e* – 7; this means that animals assume *a priori* that observations tend to be generated by a small number of causes.

Conditional on the latent context, a number of random variables are observed. In particular, on each trial a single episode of a MDP is played. (We index trials *t* and steps within each trial as *i*). The resulting state trajectory is determined, conditional on the context and the agent’s action choices, by an initial state distribution *P*(*s*_0,*t*_ = *s*|*k*_*t*_) and a state-action-state transition function *P*(*s*_*i* + 1,*t*_ = *s*|*s*_*i*,*t*_, *a*_*i*,*t*_, *k*_*t*_). These functions are defined over a set of states *s* ∈ *S* that are shared across contexts. We also assume the state-reward mapping *r*_*i*,*t*_ = *f*(*s*_*i*,*t*_) is deterministic and shared across contexts. (This is because we assume the state identity itself – e.g., illness – directly determines the state’s utility.) Alongside the MDP state trajectory, on each trial the context also emits *J* binary features, with probabilities *P*(*f*_*j*,*t*_ = 1|*k*_*t*_), meant to capture environmental features that are constant during the trial and action-independent. A priori, each feature’s probability, the initial state, and the state-action-state transition functions are each independent uniform (i.e., Beta(1,1) or Dirichlet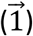).

### Inference

The goal of the learner is to learn the MDP contingencies (notably the state transition model) so as to evaluate the long-run reward consequences for candidate actions. This, in turn, requires inferring the mapping from trials to latent contexts. Following (Fearnhead, 2004; Gershman et al., 2010; Sanborn et al., 2006), we approximate inference in this model using a Rao-Blackwell particle filter to represent hypotheses about the sequence of latent context identities with an ensemble of samples. At the start of each trial *t*, each particle *m* represents a candidate assignment of all previous trials to contexts; collectively, the ensemble of particles are samples from the posterior distribution over such partitions conditioned on previous experience through trial *t* − 1. This property is maintained recursively using a combined sampling-resampling step at the end of each trial *t*, whereby a new ensemble of particles is sampled (with replacement) from all possible extensions of the previous particles plus a context assignment for trial *t* (Sanborn et al., 2006). These are sampled proportional to the prior probability of the context (from the Chinese restaurant process equations conditioned on each particle’s previous context sequence) times the likelihood of the observations during trial *t* (which is analytically computable, conditional on the particle’s previous and proposed current contexts), normalized over all particle-context combinations.

Importantly, conditioning on samples of the context assignments reduces the rest of the model learning problem (inferring the posterior distribution over the per-context feature, initial state, and state transition functions) to the same simple form as in previous theories (Daw et al., 2005; Gershman et al., 2010; Keramati et al., 2011). With the context assignments for particle *m* known, the exact posterior distributions over the per-context observation distributions each have conjugate Beta or Dirichlet forms. Updating these per-particle/per-context distributions then correspond to the standard procedure of counting the features, states, and state transitions observed in each context, according to particle *m*’s sampled context sequence. For instance, the Bernoulli probability that feature *f*_*j*_ = 1 for context *k* in particle *m* is *Beta* (1 + *N*_*j* = 1|*k*,*m*,_ 1 + *N*_*j* = 0|*k*,*m*_), where the *N*s count features observed on each visit to the context, added to the initial pseudo-counts from the *Beta*(1,1) prior. Thus, at the end of each trial, for each resampled particle, we increment feature and state counts in the context to which that particle assigned the trial, for the observed features and state trajectory.

### Action selection

Model-based planning involves using the value iteration algorithm recursively to compute the expected return for each candidate action in the current state, 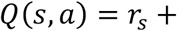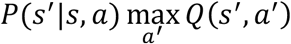 (Sutton & Barto, 2018). This in turn depends on the learned state transition function *P*(*s*′|*s*, *a*). In the usual (single, known context) case, this is usually taken as the mean of the Dirichlet posterior distribution, *P*(*s*′|*s*, *a*) ∝ 1 + *N*_*s*′|*s*, *a*’_, where *N* counts previous state transitions (and 1, is again, the initial pseudocount from the prior; e.g., Daw et al., 2005).

In the current setting, the same state and action can imply different state transition distributions in different contexts; moreover, there is uncertainty both about which context is currently active and which contexts were encountered on previous trials. We again use the ensemble of particles to marginalize all these at each choice step during trial *t*. The high-level strategy is to compute a state transition function in expectation over all this uncertainty by averaging the mean transition distribution associated with each possible context (as a candidate current context) within each particle (as a candidate assignment of previous trials to contexts), and then further averaging these transition functions over particles. Finally, we compute *Q*(*s*, *a*) for each candidate action in the current state using the resulting net transition function. Note that this procedure is approximate (for instance, it neglects correlations in state encounters over timesteps induced by the latent cause structure). Also note that although before each trial starts, all particles *m* are equally likely under the posterior (and each particle then specifies a prior probability over contexts *k*), once context features and states are observed in a trial, this additional evidence affects the conditional probability of particles and of contexts within each particle. We thus average contexts (within particles) weighted by their posterior probability, prior times likelihood, given the observations (states and features) so far in the trial, and similarly importance weight the particles by the associated marginal likelihoods of the observations. The agent selects an action softmax in the *Q* values (*P*(*a*|*s*) ∝ exp(*βQ*(*s*, *a*))); a new state is observed and the process is repeated (recomputing the importance-weighted marginal transition function and repeating value iteration for the next choice) until reaching a terminal state.

### Task

We simulate the effect of reward devaluation on instrumental conditioning using a stylized version of the task (**Figure 2**, after Daw et al., 2005, Piray et al., 2011; see Methods), which preserves the logic of sequential action-outcome evaluation while replacing self-paced free-operant leverpressing with a more structured trial-based MDP in which each trial contains a series of discrete binary choices (e.g., whether to leverpress or whether to consume a pellet). Using different series of state and outcome encounters in this setting, we also simulate the food-illness pairing taste aversion^1^ trials and unpaired illness control trials, along with the exposure experiences given to each group to counterbalance these experiences. Finally, we also conduct an instrumental extinction test and the outcome consumption test again using the same states.

## Results: persistent instrumental responding for a devalued reinforcer

The traditional empirical signature of habits is persistent instrumental responding for a devalued reinforcer (**Figure 1c**). The key insight of the current model is that such insensitivity can arise not only because the action itself is chosen by model-free or stimulus-response methods, but if the taste aversion training is inferred to arise from a distinct latent context than the leverpress training and test. In this case, even though the decision whether to leverpress is entirely model-based (that is, it is informed by anticipating and evaluating the food outcome), the taste aversion conditioning experience does not apply to these calculations, but is instead viewed as relevant only in a different context.

We first examine this phenomenon by simulating recent experiments (Bouton et al., 2021) that explicitly manipulate the *overt* context for the leverpress training vs. taste aversion conditioning (e.g., by conducting the procedures in distinct physical environments), which demonstrate that the effect of taste aversion conditioning on instrumental training is modulated by contextual similarity. Having understood the behavior of the model in this setting, we move on to consider the effect covert contextual grouping even in a single physical setting.

### Aversion Conditioning Context

First, we examined whether the physical context in which aversion conditioning took place had an influence on subsequent lever pressing, as observed recently empirically (Amaya et al., 2020; Bouton et al., 2021). To do this, we simulated the reinforcer devaluation paradigm for paired and unpaired conditions (**Figures 1a** and **1b**, **2a** and **2b**), while also varying the context in which the aversion conditioning occurred, which could either be the same as that for the instrumental acquisition and test phases or different. (In the simulations this occurs by changing the contextual features that are present or absent in each phase, see **Table 1**) The basic empirical finding (Bouton et al., 2021) is that animals in the paired condition demonstrate reduced leverpressing in extinction relative to unpaired controls; but this sensitivity of instrumental responding to reward devaluation is abolished when the leverpress training and aversion conditioning occur in different contexts. This result presumably reflects a failure to generalize the aversion to the food across contexts when deciding whether to leverpress which results in a pattern of devaluation-insensitive responding similar to that usually interpreted in terms of habits.

**Table 1.** Trial setup used to compare reward devaluation when taste aversion conditioning occurs in different contexts – aversion same (where animals undertake the taste aversion conditioning in the same context as during acquisition) and aversion different (where these phases occur in different contexts such as the animals homecage and the instrumental test chamber). Refer to **Figure 2** for the corresponding MDPs. *unpaired condition

|  | N trials | Start State | Aversion Same |  | Aversion Different |  |
| --- | --- | --- | --- | --- | --- | --- |
|  |  |  | Lever Features (n=3) | Contextual Features (n=3) | Lever Features (n=3) | Contextual Features (n=3) |
| **Acquisition** | 30 | $S_1$ | 1 | 1 | 1 | 1 |
| **Aversion Conditioning**<br>(injection + pellet delivery / pellet delivery*) | 15 | $S_2$ | 0 | 1 | 0 | 0 |
| **Aversion Conditioning**<br>(context exposure/ injection*) | 15 | $S_4$ | 0 | 1 | 0 | 0 |
| **Lever Test** | 1 | $S_1$ | 1 | 1 | 1 | 1 |
| **Consumption Test** | 1 | S<sub>2</sub> | 0 | 1 | 0 | 1 |

Similar to the pattern observed empirically (Bouton et al., 2021), simulations from our model revealed that in the lever test (**Figure 3a**) there was a significant interaction between condition (paired, unpaired) and context (same, different) (F(1, 76) = 19.81, p<0.001). This was the result of reduced lever pressing in the lever test between paired and unpaired conditions when the aversion conditioning occurred in the *same* context as the acquisition phase (t(26.71)=−6.16, p<0.001), a difference which was absent when this occurred in a *different* context (t(37.68) = – 0.17, p=0.87). As shown in **Figure 3c** and **3d**, these results reflect the model’s ability to assign different phases of training to the same or different latent causes, and thereby modulate generalization across them. In particular, aversion training is assigned to the same or a different latent cause as that for instrumental training and testing, when it occurs in the same or a different physical context respectively. This separation in the different conditions is driven by different environmental cues, and results in the decision whether to leverpress being unaffected by the aversion conditioning.

**Figure 3.**
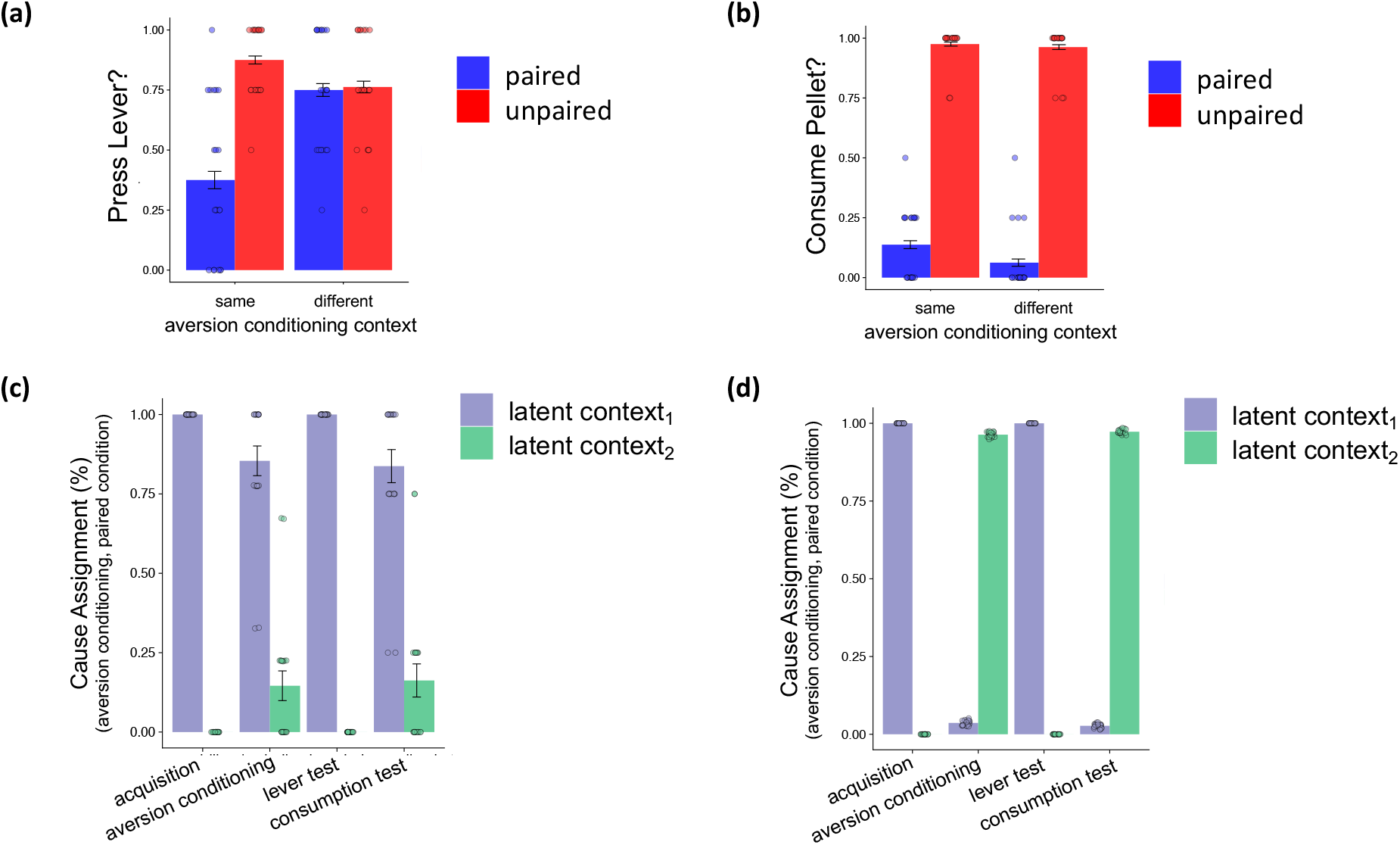
**(a)** In the lever test, simulations revealed reduced lever pressing in the lever test between paired and unpaired conditions when the aversion conditioning occurred in the *same* context as the acquisition phase (t(26.71)=−6.16, p<0.001), a difference which was absent when it occurred in a *different* context where lever pressing similar between the paired and unpaired conditions (t(37.68) = −0.17, p=0.87). **(b)** In the consumption test, agents were reluctant to approach and consume pellets regardless of where the aversion conditioning had taken place (paired vs unpaired different: t(33.08) = −24.36, p<0.001; paired vs unpaired same: t(28.22) = −22.08, p<0.001, independent sample ttest). **(c)** Examining latent context assignments at each phase of the devaluation procedure when aversion conditioning occurred in the same context as acquisition revealed that – owing to the similarity of the contextual cues – the majority of trials in each phase were assigned to the same latent context (latent context_1_). **(d)** In contrast, latent context assignments when aversion conditioning occurred in an alternate context revealed that trials during aversion conditioning and the consumption were assigned to a different latent context (latent context_2_) as for acquisition and the lever test (latent context_1_).

A subtler point concerns an additional aspect of these experiments, the outcome consumption test used to verify the efficacy of the aversion conditioning. This is a second test (performed in the instrumental training context, but without the lever available), of the animal’s willingness to consume the food (“averted reinforcer”). One might assume that reduced consumption in the paired group implies that the aversion training successfully generalized to the instrumental context. But surprisingly, Bouton et al.’s (2021) data show that consumption is reduced for the paired group, even when aversion conditioning occurred in a different context: that is the consumption and instrumental leverpressing tests are dissociated in this regard. The model also captured this result: agents were reluctant to approach and consume pellets regardless of where the aversion conditioning took place (paired vs unpaired different: t(33.08) = −24.36, p<0.001; paired vs unpaired same: t(28.22) = −22.08, p<0.001, independent sample ttest, **Figure 3b**) with no interaction between condition (paired, unpaired) and aversion conditioning context (F(1, 76) = 1.36, p=0.24). In the model (**Figure 3d**), this occurs because the consumption test tends to be assigned to the same latent cause as the aversion training rather than to the instrumental context; this in turn relates to the fact that although the environmental cues match those of the instrumental context, other aspects of the situation (notably the start state and the absence of the lever) are closer to the aversion training context.

Another consequence of this different pattern of cause assignments (**Figure 3c, d**) is that agents are slow to learn to avoid pellet consumption during aversion conditioning when this occurs in the same context as the acquisition phase (**Figure 4**). This occurs through latent inhibition as the latent context contains a history of pellet consumption leading to rewards carried over from acquisition that it needs to be overwritten via new learning in the aversion conditioning phase. In contrast, when aversion conditioning is assigned to an alternate latent cause as acquisition, agents are quicker to stop consuming the pellet as they do not need to overwrite these new experiences (consuming pellet leads to illness) with the old one (consuming pellet leads to reward). This is a similar pattern to that which has been observed empirically (Amaya et al., 2020; Bouton et al., 2021).

**Figure 4.**
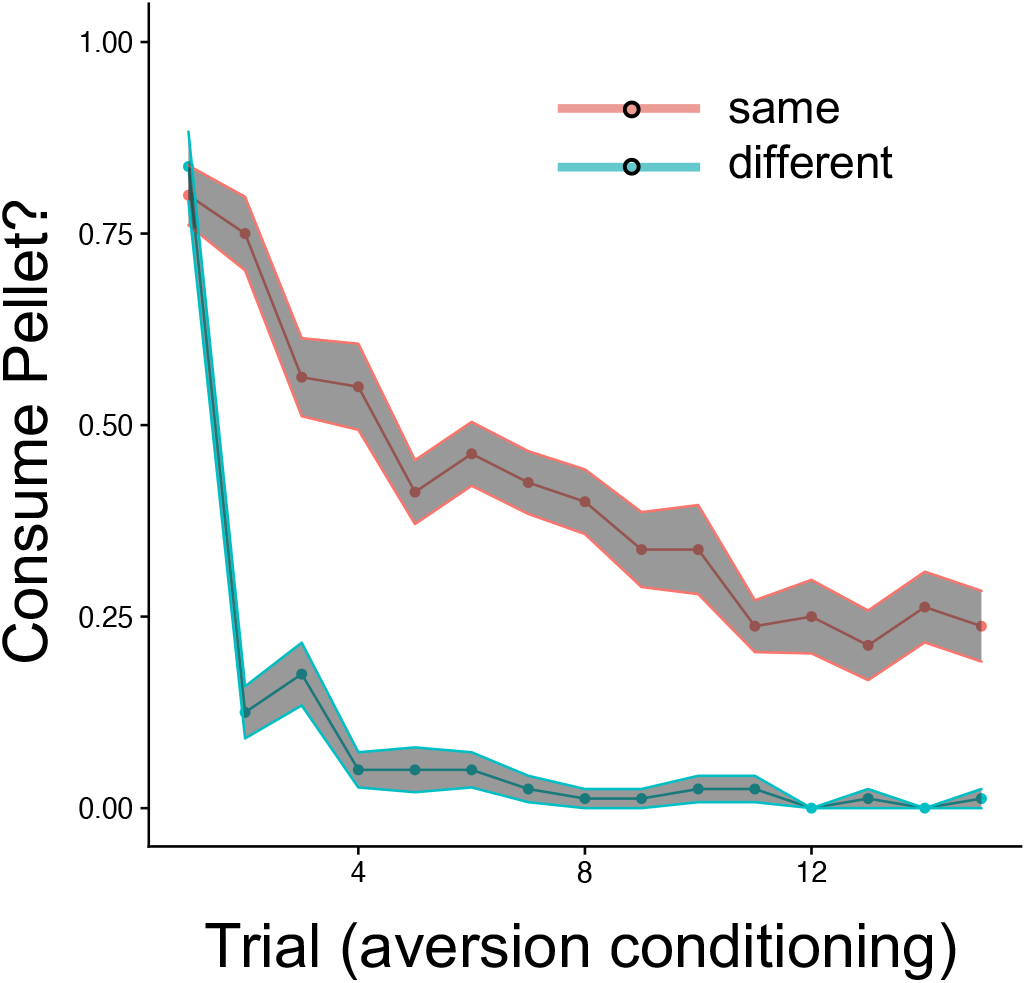
A consequence of assigning experiences during taste aversion conditioning to a new latent cause (which occurs for aversion other) is that agents can quickly learn to avoid consuming the pellet as this new latent cause has no history it needs to overwrite. In contrast, when aversion conditioning is assigned to the same latent cause as acquisition (which occurs for aversion same), agents are slower to stop consuming the pellet as they need to overwrite the new experiences (consuming pellet leads to illness) with the old one (consuming pellet leads to reward).

### Length of training

The foregoing simulations demonstrate that even though the model decides whether to leverpress using only model-based evaluation of the outcome, it can produce habit-like insensitivity of instrumental responding to reinforcer devaluation when the training context is manipulated to discourage generalization between the task phases. A question remains whether the model can also produce such ersatz habits even in the more usual experimental circumstance in which this is observed: when all training and testing occur in a single context, but the instrumental response is overtrained.

We thus examined whether length of training trials during instrumental acquisition generated differences in lever pressing post devaluation in the lever test (see **Table 2** for parameters used in the simulations). Entering mean lever pressing scores in the lever test from each simulation (N=20) into a factorial regression on length of training and condition revealed a significant interaction of these factors (F(1, 76) = 16.65, p<0.001). This was the result of a difference between paired and unpaired conditions under moderate training (t(26.71) = −6.16, p<0.001) which was absent under extended training (t(33.37) = −1.45, p=0.16; **Figure 5a**). Comparing lever pressing following extensive training relative to moderate training in the paired condition revealed lever pressing in an extinction test to be significantly greater following extensive training (t(33.44) = 4.77, p<0.001, independent sample ttest) with no difference between training regimes observed for unpaired (t(38) = −0.26, p=0.80). Again, examining responses in the consumption test (which proceeded the lever test) confirmed the efficacy of the aversion conditioning in both cases (**Figure 5c**) – agents were reluctant to approach and consume pellets in the paired condition relative to the unpaired condition both under moderate (t(28.22) = −22.08, p<0.001)) and extensive training regimes (t(28.05) = −33.27, p<0.001) however the difference was marginally greater following extensive training (condition*training interaction: F(1, 76) = 3.46, p=0.07).

**Table 2.** Trial setup used to compare moderately trained with extensively trained animals. The main difference is the number of trials during acquisition. Refer to **Figure 2** for the corresponding MDPs. *unpaired condition

|  | N trials |  | Start State | Lever Features<br>(n=3) | Contextual Features<br>(n=3) |
| --- | --- | --- | --- | --- | --- |
|  | Moderate Training | Extensive Training |  |  |  |
| **Acquisition** | 30 | 60 | S<sub>1</sub> | 1 | 1 |
| **Aversion Conditioning**<br>(injection + pellet delivery / pellet delivery*) | 15 | 15 | S<sub>2</sub> | 0 | 1 |
| **Aversion Conditioning**<br>(context exposure/ injection*) | 15 | 15 | S<sub>4</sub> | 0 | 1 |
| **Lever Test** | 1 | 1 | S<sub>1</sub> | 1 | 1 |
| **Consumption Test** | 1 | 1 | S<sub>2</sub> | 0 | 1 |

**Figure 5.**
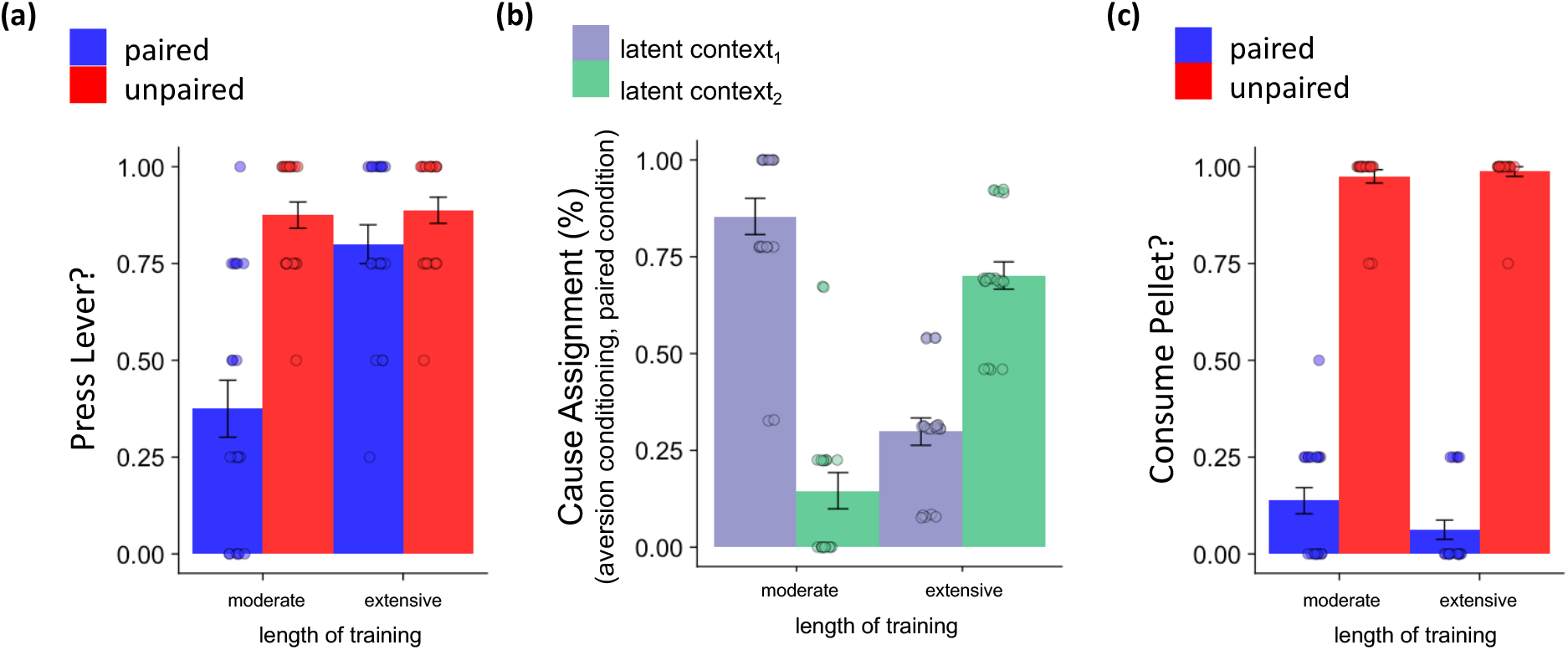
**(a)** Propensity to lever press in the lever test following moderate and extensive training. We observed an interaction between condition and training length (F(1, 76) = 16.65, p<0.001), the result of greater lever pressing following extensive training relative to moderate training in the paired condition (t(33.44) = 4.77, p<0.001, independent sample ttest) with no difference between training regimes observed in the unpaired condition (t(38) = −0.26, p=0.80). This pattern of results closely resembles the pattern observed empirically (Holland, 2004) despite arising from an exclusively model based learning system absent of habits. **(b)** Cause assignment during aversion conditioning under moderate and extensive training for the paired condition simulations. Under moderate training, the majority of transition sequences observed and used to update transition knowledge are assigned to latent context 1, the same context as experiences during acquisition. Under extensive training, the majority of transition sequences observed are assigned to latent context 2, an alternate context to the context experiences observed during acquisition are assigned to. **(c)** In the consumption test, agents were reluctant to approach and consume pellets regardless of the length of the acquisition training (paired vs unpaired moderate: t(28.22) = −22.08, p<0.001; paired vs unpaired extensive: t(28.05) = −33.27, p<0.001, independent sample ttest). Bars represent averages over the simulations (N=20). Individual data points represent the average for each simulations. Error bars represent standard error of the mean.

This pattern of results – greater lever pressing in an extinction test under extended relative to moderate training – closely resembles the pattern observed empirically (Adams and Dickinson, 1981). However, whilst this phenomenon has in the past been interpreted as evidence of habit formation in the case of extended training, here it emerges in the service of an exclusively model based learning system absent of habits (i.e., without a model free learning component). To understand how these differences in lever pressing could arise out of a purely model based system, next we examined the cause assignment during the aversion condition phase in our simulations. This revealed that following moderate training, the majority of experience accrued during aversion conditioning was assigned to the same latent context as the context experiences during acquisition were assigned to (latent context 1, **Figure 5b**). In practice this has the effect that experiences learnt during acquisition – specifically that consumption of a pellet leads to reward – are overwritten during aversion conditioning – as consumption of a pellet now leads to a loss (illness). Following extensive training however, more aversion conditioning experiences are assigned to a separate latent cause (latent cause 2) to those that acquisition learning experiences are assigned to. In practice, this has the effect that state action state sequences learnt during acquisition are “protected” from new experiences encountered during aversion conditioning. Hence the animal does learn that pellet consumption leads to illness but segments this experience to an alternate transition model, leaving the original transition model learnt in acquisition intact. The change over training in the tendency to lump vs. split experiences across causes reflects a characteristic feature of statistical inference in this type of model: the model starts with a bias to favor simpler models (the so-called Bayesian Occam’s razor), and it takes additional evidence to justify a more complex model containing multiple latent causes (Courville et al., 2003).

## Discussion

Computational models have separately explored two distinct mechanisms relevant to conditioning. The first concerns the strategy for evaluating the decision variable. The fact that instrumental responses are sometimes sensitive – but sometimes insensitive – to reinforcer devaluation has been argued to reflect the use of two learning mechanisms, a deliberative model-based mode of control and an automatic model-free mode of control (Daw et al., 2011, 2005). In the terms of traditional associative learning models, these findings are, analogously, interpreted in terms of goal-directed action-outcome vs habitual stimulus-response responding (Dickinson, 1985; Dickinson and Balleine, 2002). However, a second mechanism – heretofore studied mostly in the context of Pavlovian conditioning – concerns how animals track different contingencies in different situations, by grouping experiences into different covert latent causes (Gershman et al., 2010). This grouping process leads to differential generalization across them.

Here, we present a new model combining elements of both lines, to explore the consequences of latent cause inference for instrumental conditioning. We show that an exclusively model-based learner can show insensitivity to reinforcer devaluation due to a failure to generalize taste aversion conditioning to the lever test. The model reproduces patterns of behaviors previously thought to be a signature of habits: persistent instrumental responding in extinction for an averted reinforcer. We show this can arise following extended training (Adams, 1982) and when the aversion conditioning and acquisition/test contexts differ (Bouton et al., 2021). In both cases, the model captures value-insensitive responding within each cause due to failure to generalize aspects of taste aversion learning to the latent cause where the instrumental test is inferred to occur. Taste aversion experience generalizes more readily during the consumption test because of greater feature overlap (e.g., the absence of the lever). Although these results do not rule out a contribution of model-free learning, they point to the importance of state inference in instrumental learning.

One influential application of habit models has been as a candidate substrate for seemingly consequence-insensitive choices in compulsive disorders including drug abuse (Everitt and Robbins, 2016, 2005). The present work offers a new interpretation of this view, and specifically a different perspective on experimental evidence (much of it from instrumental devaluation and related procedures). In general, the specificity of learning to a latent context – and the failure to generalize that learning across contexts – offers another way of explaining why behaviors may be detached from their consequences (potentially to deleterious ends), even when actions are chosen in a fully deliberative, model-based manner. Indeed, one common criticism of the habit view of addiction is that although it can explain some well-trained stereotyped actions (e.g., drug consumption) it seems difficult to explain many behaviors involved in drug-seeking, which seem to involve outcome-specific deliberation (e.g., craving) and seemingly model-based or goal-directed ability to choose novel actions (Daw, 2015; Tiffany, 1990). The emphasis on the contextual specificity of learning in the current model (and latent cause models generally) also connects naturally to much data on contextual sensitivity in drug abuse, including relapse and craving (Bossert et al., 2004; Bouton and Swartzentruber, 1991; Crombag and Shaham, 2002; Wikler, 1973).

There is also more specific evidence leveraging the reinforcer devaluation procedure to investigate a putative habitual basis for drug seeking behavior (Clemens et al., 2014; Corbit et al., 2012; Dickinson et al., 2002; Miles et al., 2003) generally reporting that drug reinforcers support habitual (devaluation insensitive) instrumental responding more readily than natural ones. An alternative interpretation of some of these results under the current model is that (in addition to or instead of promoting model-free responses) drug reinforcers promote a greater tendency to split causes and fail to generalize between them, for instance because of the salience of drug cues or enhanced salience attributed to other cues in the presence of drugs (Field and Cox, 2008), and/or because the acute intoxicating effects of the drug itself serve as an additional context cue. In humans, drug addiction has also been associated with reduced model-based behavior on a “two-step” Markov decision task (Gillan et al., 2016; Voon et al., 2015) whose logic is similar to reward devaluation. In the current model, such behavior might again alternatively reflect a greater tendency in these disorders to group a subset of trials (“rare transitions”) into a distinct latent cause. Overall, without ruling out a contribution of model-free or habitual processes to drug abuse, the present model offers an additional potentially contributing mechanism, and also may explain aspects of drug abuse (such as craving and contextual sensitivity) not easily understood by habit accounts.

Although we eschew model-free learning altogether in the current model in order to emphasize the effect of latent cause inference even on model-based learning, we think it quite likely that the brain also employs model-free learning, and devaluation-insensitive instrumental leverpressing is therefore multiply determined. For instance, the current work explains many phenomena of habits when devaluation occurs via taste aversion conditioning, but an alternative approach (which produces broadly similar results) instead manipulates the animal’s motivational state, studying instrumental responding under satiety (Balleine, 1992). In general, these results appear less easily explained by latent cause inference, because the motivational state itself would enter into the contextual inference in a way that would tend in any case to discourage instrumental responding. In any case, future work could consider the integration of model-free learning back into the current model (see also Schwober, 2021); in which case the parsing of experiences among causes would be expected to affect the progression of learning from model-based to model-free within each cause (Daw et al., 2005) whereas switching across causes might drive unlearning and renewal of habits (Miller et al., 2019; Schwöbel et al., 2021; Smith et al., 2012). Finally, other recent research has emphasized other optimizations or simplifications of model-based choice short of fully model-free habits, including temporal abstraction (Russek et al., 2017), pruning (Huys et al., 2012; Mattar and Daw, 2018) and model sharing (Glitz et al., 2022), all of which might potentially interact with latent cause inference in an extension of the current work.

Nonetheless, the results we present here show that by integrating beliefs about state dynamics into a latent cause inference model the breadth of behaviors that can potentially be accounted for under a purely model-based learner is larger than previously appreciated.

## Methods

### Simulations

First, we simulated 30 trials of instrumental acquisition 20 times, varying the context in which the taste aversion conditioning phase occurred (by altering the contextual cues present/absent, see **Table 1**). In one set of simulations (*aversion same*) the taste aversion conditioning occurred in the same context as the acquisition context. In another (*aversion different*) the taste aversion conditioning occurred in a different context.

Next, we simulated each condition (paired, unpaired) 20 times for two different training lengths: *moderate training* – in which the acquisition phase lasted for 30 trials – and *extensive training* – in which the acquisition phase lasted for 60 trials (double the number of trials as the aversion conditioning phase). In these simulations, the contextual features present/absent in each phase were matched between conditions (paired/unpaired) and training durations (moderate/extensive, see **Table 2**).

### Paired Condition

The MDPs used in each simulation for each condition are displayed in **Figure 2**. The contingency between these states, actions and subsequent states changed between phases. In the paired condition (**Figure 2a**), during the Acquisition phase animals began in S_1_ and selected whether to press a lever or not. Pressing a lever delivered a pellet (transition to S_2_) where they faced a second action choice: whether to consume the pellet or not. Consumption of the pellet transitioned to a terminating state with a positive reward (S_5_). Decision not to consume the pellet (from S_2_) or not to press the lever (from S_1_) terminated the episode without a reward (transition to S_3_). The Taste Aversion Conditioning phase was separated into two sections. In the first section, agents began in S_2_ where a decision to consume a pellet now transitioned to a terminating state with a negative reward (transition to S_6_). In the second section, animals began in an “context exposure state” (S_4_) where an obligatory action transitioned them to a terminating state with no reward (S_6_). This was done such that animals had both the same amount of “pellet consume” decisions and exposure to the aversion conditioning context in each condition (Bouton et al., 2021). Lever Test and Consumption Test phases each consisted of a single trial and were exactly as described for Acquisition and Taste Aversion Conditioning (section 1) phases respectively except that a decision to lever press (from S_1_ in the Lever Test) now led to S_3_ (to mimic the fact that lever presses in the test phases are usually carried out in extinction).

### Unpaired Condition

Start states and transition dynamics for the unpaired condition (**Figure 2b**) were the same as the paired condition for the Acquisition and two test phases. However the taste aversion conditioning phase differed. This phase was again separated into two sections. In the first section, animals began in S_2_ where a decision to consume a pellet continued to transition to a terminating state which accrued a positive reward. [Note this is different to the paired condition where consumption of a pellet in this phase led the agent to a terminating state with a negative reward.] In the second section, animals began in an “injection state” (S_4_) where an obligatory action transitioned them to a terminating state with negative reward (S_6_). This therefore (in principle) unpairs transitions to the negative reward state (i.e. illness) and pellet consumption.

In each simulation we ran 4 virtual agents (animals). “Lever press” actions in the lever test and “consume pellet” actions in the consumption test were averaged per simulation (i.e., over the 4 agents) and then these per simulation scores were averaged (over the 20 simulations) to get an overall mean tendency to lever press and pellet consume. Code and simulations were run in MATLAB (2020b). In all simulations we used set the concentration, *α*, equal to 1*e* – 7. The slope of the softmax function, *β* was drawn from a normal distribution on each trial with a mean of 6 and a standard deviation of 1. We used a particle filter with 3,000 particles. We set the maximum number of causes in the model to 10 (although in principle generative models allow for an ever-expanding number of latent causes, setting a low value of *α*, as we do here, meant that in practice the number of latent causes established at the end of each simulation never exceeded 3).

## Acknowledgements

This research was supported by a US Army Research Office grant (W911NF-16-1-0474) to N.D.D. and a Sir Henry Wellcome Postdoctoral Fellowship (209108/Z/17/Z) to N.G. We would like to thank Mark Bouton for helpful insights and discussions.

## Footnotes

1 We use the terms aversion conditioning/trials/training in parts of the manuscript as an abbreviation of the more complete terms *taste* aversion conditioning/trials/training.

## Notes

### Competing Interest Statement

The authors have declared no competing interest.

